# Engineered isoprene production from *Chlamydomonas reinhardtii* using herbicide selection markers and CO_2_-fed cultivation optimization through multi-parallel photobioreactor headspace gas analysis

**DOI:** 10.1101/2025.04.22.649625

**Authors:** Razan Z. Yahya, Sebastian Overmans, Gordon B. Wellman, Kyle J. Lauersen

## Abstract

Metabolic engineering requires selection markers for transformant generation. However, use of antibiotic resistance is of concern for potential horizontal gene transfer in the environment. Herbicide resistance markers are an alternative for photosynthetic cell line engineering as these agents are plant-specific with resistance mechanisms that can be generated from mutations of endogenous genes. Here, we developed norflurazon and oxyfluorfen resistance markers for nuclear genome transformant selection in the model green alga *Chlamydomonas reinhardtii*. These were used to engineer robust isoprene biosynthesis by facilitating overexpression of a yeast isopentenyl-diphosphate delta-isomerase (*Sc*IDI), the alga’s own beta carotene ketolase (*Cr*BKT), and the sweet potato isoprene synthase (*Ib*IspS). Further UV-C mutagenesis and colony selection were employed to improve yields to ∼350 mg isoprene L^-1^ culture on organic carbon. It was then possible to optimize CO₂-driven cultivation and isoprene biosynthesis in batch and continuous processes using multi-port, real-time, in-line mass spectrometry coupled to parallel photobioreactors. The highest isoprene yields in batch were achieved under 900 µE illumination and 33 °C and, in turbidostat mode, ∼51 mg isoprene L culture^-1^ day^-1^ was achieved for 3 days concomitant with algal biomass production. Cultivation of the engineered alga directly in effluent from an anaerobic membrane bioreactor was also conducted. Isoprene production was concomitant with removal of ammonium and phosphate from the wastewater, and biomass production was similar to that in replete medium. Isoprene yields exhibited gradual reduction after each successive repetitive refresh, which could be mitigated by supplementation of trace elements. The results demonstrate that engineered algae could be used as a secondary wastewater treatment step while generating both biomass and volatile co-products like isoprene.

**Graphical Abstract:** 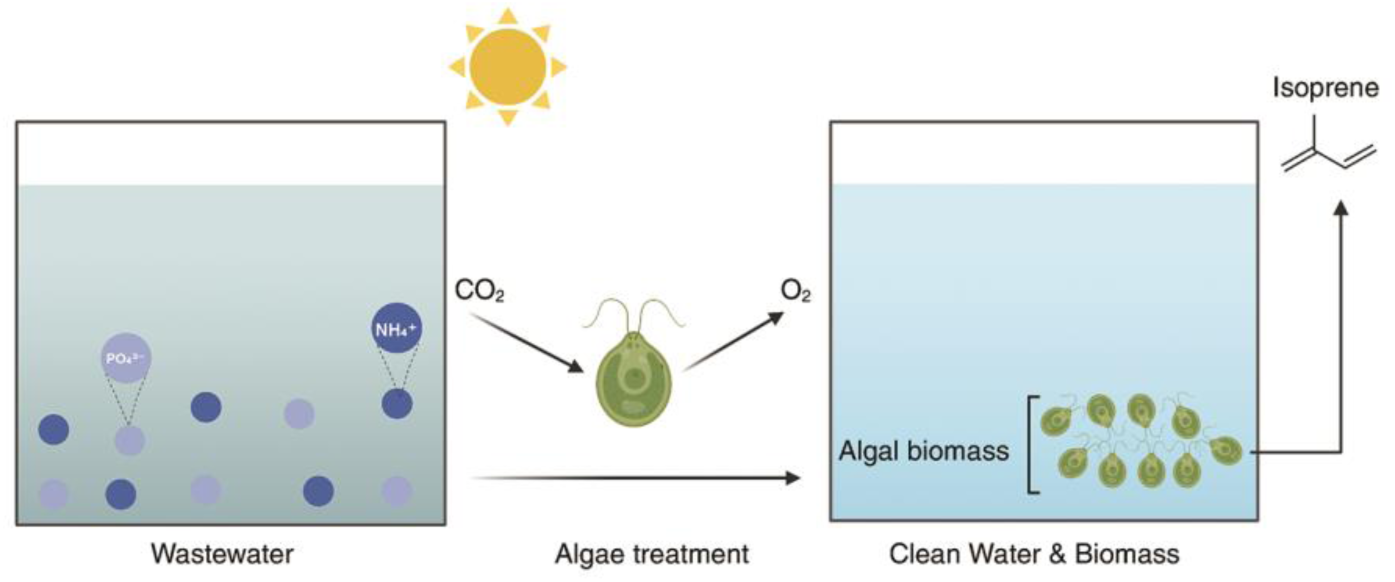

## 1 Introduction

As photosynthetic microbes, algae have been proposed as sustainable systems for the conversion of inorganic chemicals, like carbon dioxide, ammonium, nitrate, and phosphate into their biomass, which is rich in protein, carbohydrates, lipids and pigments (Goold et al., 2024; Janssen et al., 2022; Schipper et al., 2022; Torres-Romero et al., 2019; Wichmann et al., 2020). These inorganics are often found in human activity effluents and emissions, which position algae as waste stream-conversion vehicles if their cultivation is coupled to point sources (Blifernez-Klassen et al., 2023; Goswami et al., 2022; Henkanatte-Gedera et al., 2017; Nguyen et al., 2022; Selvaratnam et al., 2015; Silkina et al., 2019). Algal cultures can be installed on any land-use type, even directly coupled to industrial sites (Hallmann, 2016; Posten, 2009). Their cultivations can yield clean water and algal biomass, which has numerous applications from animal feed to specialty chemical production. Genetic engineering of algae has been proposed as a means to add layers of value to algal bioprocesses by yielding novel pigments, specialty or platform chemicals, and even volatile co-products that can be harvested concomitantly (Einhaus et al., 2023; Goold et al., 2024; Lauersen, 2019; Wichmann et al., 2020).

The 5-carbon hemiterpene isoprene is one such volatile chemical which can be produced from engineered algae (Yahya et al., 2023). Isoprene is the monomer of rubber and a platform chemical used in synthesis applications (Matos et al., 2013). Biotechnological production of isoprene is possible through the expression of plant isoprenoid synthases in microbial hosts (Miller et al., 2001; Whited et al., 2010). Although not yet commercially feasible, producing isoprene from engineered photosynthetic microbes presents the possibility of generating isoprene from carbon dioxide (Chaves & Melis, 2018; Matos et al., 2013). This strategy is even more attractive if the nutrients for algal growth, nitrogen and phosphorous, come from wastewater, thereby generating a useful chemical that can be harvested from the gas phase of a culture separately from its biomass (de Freitas et al., 2023; Yahya et al., 2023).

*Chlamydomonas reinhardtii* has become a model for algal metabolic engineering aimed at additional chemical and bioproduct generation (Jackson et al., 2021; Perozeni & Baier, 2023; Schroda, 2019). Transformation of its nuclear genome is straightforward, and auxotrophic complementation has been established for 35 years (Kindle, 1990; Kindle et al., 1989). However, overexpression of heterologous transgenes from this genome has historically been challenging and only recently normalized in practice through decades of trial and error-built understanding of its nuanced transgene design requirements and genetic regulation (Reviewed in (Perozeni & Baier, 2023)). Integration of foreign DNA into the nuclear genome occurs primarily through non-homologous end-joining (NHEJ) (Gumpel et al., 1994), and transgene cassettes are also randomly cut by nucleases *en route* to the genome (Gutiérrez et al., 2022). These phenomena hinder efficient transformation of large, multigene constructs, necessitating serial transformations and selection of transformants expressing genes of interest in a stepwise manner, usually employing multiple antibiotic resistance markers (Amendola et al., 2023; Einhaus et al., 2022; Freudenberg et al., 2022; Gutiérrez et al., 2024; Lauersen et al., 2018; Zhao et al., 2023). Generating algae with multiple antibiotic resistance genes increases risks of their horizontal gene transfer to the environment and limits a strain’s use in scaled cultivations, especially outdoors, or those aimed at waste-recovery and/or water treatment. Our recent report demonstrated that engineered *C. reinhardtii* could produce volatile isoprene as a co-product to its biomass (Yahya et al., 2023). The strains produced up to ∼360 mg L^-1^ isoprene in headspace concomitant with algal biomass in culture. However, these strains contain three antibiotic resistance markers. We, therefore, sought alternatives that could be more appropriate for scaled algal cultivations and waste-stream conversion efforts.

Previous reports have shown that herbicides can be effective selection agents for *C. reinhardtii*, including the protoporphyrinogen oxidase (PPO, ‘protox’) inhibitor oxyfluorfen in addition to norflurazon, which targets the enzyme phytoene desaturase (PDS) (Boger et al., 1998; Bruggeman et al., 2014; Randolph-Anderson et al., 1998). Oxyfluorfen inhibits *C. reinhardtii* at nM concentrations, making it particularly suitable for large-scale liquid algal cultures. Resistance to each herbicide is mediated by mutations in endogenous genes, protox or PDS (Randolph-Anderson et al., 1998) (Bruggeman et al., 2014) (Boger et al., 1998). The bacterial PDS gene (CRTI) from *Pantotea ananatis* is naturally resistant to norflurazon and has also been used as a selection marker for *C. reinhardtii* transformation (Molina-Márquez et al., 2019). In this work, we explore the use of these herbicides *in lieu* of antibiotics to engineer *C. reinhardtii* to produce volatile isoprene.

Our previous strain engineering and isoprene quantification was conducted exclusively in small-scale organic-carbon-fed conditions using cultivation of transformants in sealed headspace vials to enable direct analysis of volatile isoprene in gas chromatography-mass spectrometry. Algal biotechnology aims to leverage the innate photosynthetic metabolism of the alga to cultivate strains on CO_2_ as a carbon source. Therefore, it was necessary to explore the capacity of our engineered cell lines to generate isoprene in light-driven CO_2_-fed cultures. For the first time for *C. reinhardtii*, we employed parallel photobioreactors coupled to in-line headspace analysis, which enabled real-time monitoring of volatile isoprene production from cultures grown on CO_2_ as a sole carbon source. Batch cultivations in 35 different light and temperature parameters identified the optimum abiotic conditions for algal growth and isoprene production. These findings were then used to run algal cultivations in both turbidostat and repetitive batch cultivations using anaerobic membrane bioreactor effluent as a medium for growth. The results presented here are a first of their kind for eukaryotic algal biotechnology and show the possibility of conducting waste revalorization concepts that generate engineered algal biomass with novel traits, concomitant separate volatile co-products, and yield clean water.

## 2 Materials and Methods

### 2.1 Algal culture conditions

The *C. reinhardtii* strain UPN22 was used as the parental strain in this study (Abdallah et al., 2022). Cultivation of *C. reinhardtii* followed the same protocol outlined in Yahya et al. (2023). Briefly, the cells were grown on TAPhi-NO_3_ medium agar plates or in 50 mL liquid TAPhi-NO_3_ medium shaken at 120 rpm unless otherwise indicated under continuous 150 µmol photons m^−2^ s^−1^ (unit referred to hereafter as µE) illumination.

### 2.2 Herbicide toxicity tests

To assess the toxicity of the herbicides norflurazon and oxyfluorfen on the parental strain *C. reinhardtii* UPN22, experiments were conducted using various concentrations of these chemicals based on a previous report (Bruggeman et al., 2014). For norflurazon, the concentrations tested were 1.5, 2.5, and 3.5 µM in liquid medium and 2, 3, and 4 µM in agar solidified medium. For oxyfluorfen, the concentrations tested were 0.07, 0.08, 0.09, and 0.11 µM in liquid medium and 0.03, 0.06, 0.09, 0.11, and 0.15 µM in agar solidified medium. TAPhi-NO_3_ medium (containing acetic acid) and TPhi-NO_3_ medium (lacking acetic acid) were used in these experiments. The cells were initially grown in 24-well plates with 1mL of TAPhi-NO_3_ medium until the population reached the late logarithmic growth phase (∼1x10^7^ cells/mL). Subsequently, cells were spun at 1000 x g for 4 min, and the pellet was washed with 700 µL of either TAPhi-NO_3_ or TPhi-NO_3_ medium. Subsequently, 100 µL of the cell suspension was added to each well of a 6-well plate in three biological replicates. Cells in TAPhi-NO_3_ and TPhi-NO_3_ media were both grown under constant 100 µE illumination for 7 d. However, cells in TPhi-NO_3_ medium were kept in an acrylic box supplied with 5% CO_2_ as the sole carbon source instead of acetic acid. In liquid culture, cell density was measured on days 1, 3, 6, and 7 using an Invitrogen Attune NxT flow cytometer (Thermo Fisher Scientific, UK), following a previously described method (Overmans & Lauersen, 2022).

### 2.3 Plasmid design, transformation and screening

Previously designed transgenes were used here for the *Ipomoea batatas* (sweet potato) isoprene synthase (*Ib*IspS), the alga’s own reengineered beta carotene ketolase (*Cr*BKT), and *Saccharomyces cerevisiae* isopentenyl-diphosphate delta-isomerase (*Sc*IDI) (Yahya et al., 2023). The *C. reinhardtii* carotenoid hydroxylase *Cr*CHYB (Cre04.g215050.t1.2) was graciously provided by Dr. Thomas Baier as previously used (Amendola et al., 2023). Other transgenes designed for expression were adapted for compatibility with the *C. reinhardtii* nuclear genome by back-translating amino acid sequences to align with the most frequent codon usage in the alga’s nuclear genome and with spread introns as previously described (Baier et al., 2018, 2020; Jaeger et al., 2019). Nuclear genome transformation was carried out as described in Kindle (1990) and outlined in (Yahya et al., 2023). Codon optimized transgenes for expression of the *P. ananatis* CrtI (Genbank: AHG94990.1), *Cr*PDS (Cre12.g509650.t1.2), and *Cr*PPO (Cre09.g396300.t1.2) variants were synthesized by Genscript (Piscataway, NJ). Cloning of variant plasmids where cassettes are mixed and matched was conducted with the pOptimized designed *Sca*I-*Xba*I or *Sca*I-*Spe*I restriction endonuclease sites (Gutiérrez et al., 2022). The final annotated sequences of all expression plasmids are provided in **Supplemental File 1**. Additionally, a detailed summary is presented in **Supplemental File 2 Tab ‘Plasmids’**. All plasmids were maintained in *Escherichia coli* DH5α cells with ampicillin as a selection antibiotic. DNA was prepared from bacterial cultures for algal transformation, as previously described (Yahya et al., 2023).

Algal Transformants were generated on agar plates with 2 µM norflurazon and/or 0.15 µM oxyfluorfen after 7 d under 100 µE for norflurazon plates and 400 µE for oxyfluorfen. The transformation efficiency was calculated as number of transformants per pmol DNA used from three separate transformation events as biological replicates. Transformants were screened for semi-quantitative assessment of gene-of-interest-fluorescent protein (GOI-FP) fusion expression following previously described methods (Gutiérrez et al., 2022). Transformants showing high target FP fluorescence, indicating high transgene expression, were selected. The correct molecular mass of each protein was verified through SDS-PAGE in-gel fluorescence of the GOI-FP fusion. For the *Cr*BKT and *Cr*CHYB transformants, cells were chosen based on a color change from green to brown or orange due to the accumulation of ketocarotenoids (Amendola et al., 2023; Perozeni et al., 2020).

2.4 Quantification of isoprene production by *C. reinhardtii*

Following the method outlined by Yahya et al. (2023), selected transformants containing all of the expected genes were inoculated from agar plates into 1 mL of TAPhi-NO_3_ liquid medium into individual wells of a 24-well microtiter plate and left shaking at 190 rpm under 100 μE for 3 d. Subsequently, 1 mL culture was transferred into 6-well plates, with 4 mL of TAPhi-NO_3_ medium, and shaken at 120 rpm until populations reached the late logarithmic growth phase. For isoprene quantification, 1 mL of each dense culture was added to 9 mL of the same medium in two biological replicates of 20 mL autoclaved headspace bevel-top vials with PTFE/silicon septum lids (Supelco, Cat.: 27306), which were immediately sealed. The vials were kept upright and subjected to continuous 100 µE for 6 d. On day 6, 0.5 mL liquid from each vial was taken through the vial septum by using a 3 mL syringe and inverting the vials for sampling with a needle to measure algal cell concentrations immediately prior to Isoprene quantification by headspace gas chromatography-mass spectrometry and flame ionization detection (HS-GC-MS-FID) following a previously described method (Yahya et al., 2023).

### 2.4 UV-C mutagenesis

The highest isoprene-producing transformants from each plasmid were subjected to UV-C mutagenesis on agar plates to generate variant colonies. For every test, liquid cultures were normalized to 1×10^6^ cells mL^−1^ and 6 mL added to a 35 mm petri dish without agar. Three different UV-C exposure times were tested on strain UPN22: 15, 30, and 60 s compared to control (0 s) by placing each plate inside a disinfection UV light box equipped with an 8-watt, 274 nm UV-C lamp (Dinies - ELG100S). After exposure, dishes were removed and the cultures shaken at 120 rpm in the dark for 24 h. Afterward, cells were centrifuged at 1000 × *g* for 4 min, the pellet was washed with 700 µL of TAPhi-NO3 medium and then plated on TAPhi-NO3 agar medium. For transformants, either one or both herbicides were added to agar plates as described above. The plates were kept in the dark for 24 h before being moved to constant light at 100 or 400 µE for 7 d to recover.

Colonies that recovered were picked up into TAPhi-NO3 agar plates containing one or both herbicides and left to recover for 2 d. Mutants were screened by fluorescent imaging as described above. Mutants exhibiting high expression signals of the desired transgenes, observed as fluorescence from GOI-reporter fusions, were selected for further analysis by SDS-PAGE in-gel fluorescence and isoprene yield as above.

### 2.5 Photobioreactor inoculation

Two strains were used in the photobioreactor experiments: the highest isoprene-producing transformant carrying the combination of plasmids 16 (*Ib*IspS-RFP-*Cr*PDS) & 13 (*Sc*IDI-mTFP- *Cr*BKT-*Cr*Protox) and its corresponding UV-mutagenized derivative. Liquid precultures were taken from selective TAPhi-NO₃ medium plates to liquid culture without herbicides. Prior to bioreactor flask inoculation, cell densities were normalized and transferred to 50 mL medium in 400 mL conical flasks. From each preculture, 50 mL was centrifuged at 1000 x *g* for 4 min. After centrifugation, the pellets from each tube were washed with fresh medium, centrifuged again and resuspended in the final desired medium. All biological replicates were from the same resuspension, and 400 mL of this low-density algal culture was transferred into each 1.5 L conical flask (Algenuity, UK). To ensure consistency across all conditions, the starting cell concentration was normalized using a flow cytometer before preparing the final culture. Growth of the algal cultures was monitored by measuring the optical density at 740 nm (OD_740_) using the built-in sensor of the photobioreactors (Algenuity, UK) throughout the experiments. OD readings were collected in 10 min intervals. All cultures were continuously sparged with a mixture of 3% CO₂ in air at a flow rate of 25 mL/min, and the pH of the cultures was continuously monitored using a pH probe.

### 2.6 Isoprene quantification using an in-line headspace analyser system

Volatile isoprene production was monitored for all experiments using a multi-port inlet real-time headspace gas analysis system coupled to the off-gas of each bioreactor, which included a triple filter mass spectrometer (Hiden Analytical HPR-20 R&D, UK) as previously described and illustrated in (Villegas-Valencia et al., 2025). The off-gas from each reactor was routed through separate gas lines, first into a 250 mL bottle containing 80 g of CaCl₂ as a desiccant. The gas was then directed to the 20-port inlet of the headspace gas analysis system. The gas composition in the headspace of each flask was analysed, with the system rotating to the next gas stream approximately every 3.5–40 minutes, depending on the number of flasks. Isoprene levels were quantified by monitoring atomic mass units (amu) 67, 68, 53, 39, 40, 41, and 27, while any overlapping amu from other gases (O₂, CO₂, N₂, Ar) were automatically deconvoluted by the Hiden Analytical QGA Software (version 2.0.8). The isoprene concentration was initially measured in parts per million (ppm) by the instrument. Additionally, the isoprene detected in each culture’s headspace was integrated over time to provide a cumulative measure of production, which was expressed as mg isoprene L^−1^ of culture. This approach ensured that both instantaneous concentration and total isoprene output over the course of the experiment were accurately captured. The output (in ppm) reported by the instrument was corrected using a standard curve **(Supplemental Figure 1)** generated from flasks containing various concentrations of pure isoprene standard, which were also kept in the Algem photobioreactors with the same gas flow. In total, 10 control flasks were established, covering five concentrations (10, 5, 2.5, 1.25, and 0.625 mg of isoprene) in duplicate. The temperature of the standard flasks was gradually raised from 20 °C to 38 °C over 1 h and maintained for 10 h, during which the isoprene standard in the headspace was quantified. The integrated isoprene amounts were plotted against the amounts added to each flask to create the standard curve. The Hiden Analytical QGA Software facilitated real-time gas analysis throughout the algal cultivations.

### 2.7 Turbidostat operation of photobioreactor

The culture was prepared as described previously, then acclimated to 33 °C and a light intensity of 900 µE for 2 d prior to the experiment. First, the culture was grown in T2Phi-NO_3_ medium at 33 °C with 900 μmol m⁻² s⁻¹ intensity and shaken at 100 rpm. The starting OD was set to 1.0, and once the OD reached 3.0, a 2 L bottle of the same medium was connected to the bioreactor via the turbidostat pump module (Algenuity, UK) **(Figure 3C)**. A standard curve was generated for the turbidostat pump flow at selected speeds prior to culture inoculation, which enabled setting the pump rpm to achieve a flow rate of 2 mL/min during turbidostat operation **(Supplemental Figure 2)**. As the OD was measured every 5 min, the pump replaced 10 mL of medium per on-cycle, equating to 2.5% of the total culture volume. The experiment continued for 5 d and isoprene production was monitored throughout the experiment using the HS analyser instrument described above.

### 2.8 Repeated batch cultivations

To test the feasibility of cultivating *C. reinhardtii* on wastewater (WW), we compared the growth performance and isoprene production of a culture maintained on replete T2Phi-NO_3_ medium against cultures grown on anaerobic membrane bioreactor (AnMBR) effluent. In total, four different wastewater treatments were used: raw effluent (‘WW’), effluent + trace elements (‘WW+trace nutrients’), or effluent with addition of either replete ammonium or phosphite according to the T2Phi media recipe (7.48 mM ammonium (NH₄⁺), 0.96 mM phosphite (PO_3_ ^3-^)). All *C. reinhardtii* cultures were inoculated and grown in Algem photobioreactors as described above, following the same preculture and washing routine with WW. One flask was prepared for each of the five treatments, and all cultures were grown for 8 d at 33°C with 900 µE intensity and shaken at 100 rpm. The pH of the cultures was monitored continuously using a pH probe, while growth of the algal cultures was monitored by measuring their optical density (OD_740_) in 10 min intervals. Optical density values were converted to biomass (g/L) using a standard curve of OD vs biomass (**Supplemental File 2**). All cultures were continuously sparged with a mixture of 3% CO₂ in air at a flow rate of 25 mL/min, and the concentration of volatile isoprene present in the off-gas (in ppb) was quantified using a multi-port inlet real-time headspace gas analysis system (Hiden Analytical HPR-20 R&D, UK), as described above. Every 48 h, 350 mL of culture was removed and replaced with fresh medium from each of the five flasks according to the respective treatment. Immediately before and after dilution, 10 mL of culture was sampled, centrifuged, the supernatant filtered, and analysed for concentration of nitrogen and phosphorus as described below.

### 2.9 Nutrient analysis

Nitrate, ammonium, and phosphorus concentrations in the culture medium were analysed using a DR 1900 spectrophotometer after preparation of the samples with LCK339, AmVer High Range, and TNT 844 analysis kits (Hach, Loveland, Colorado, USA), respectively, according to the manufacturer’s instructions. All nutrient measurements were performed in technical triplicates (*n* = 3). Briefly, 10 mL of algae culture was sampled, centrifuged (4500 x g, 5 min, 20°C), and the supernatants passed through 0.2 μm syringe filters to remove any insoluble particles before analysis.

### 2.10 Data processing

Data handling and visualization were performed using JMP Pro 18.0.2 and Python 3.11. For correlation plots, JMP was used to analyse the relationship between variables.

## 3 Results and Discussion

### 3.1 *C. reinhardtii* is sensitive to low concentrations of norflurazon and oxyfluorfen

The sensitivity of *C. reinhardtii* UPN22 to different concentrations of norflurazon and oxyfluorfen was tested to identify optimal selection conditions for transformation on both agar plates and in liquid medium. Norflurazon was effective in both agar plates (2 µM) and liquid medium (1.5–3.5 µM), resulting in near-complete growth inhibition by day 7 in liquid cultures **(Figure 1A)**. Oxyfluorfen exhibited a more gradual, dose-dependent effect on agar plates (0.03–0.15 µM), with complete growth inhibition observed at 0.15 µM, but had no significant effect on liquid cultures at concentrations of 0.07–0.11 µM **(Figure 1B)**. Both organic carbon- and CO_2_-grown cultures were tested, with oxyfluorfen on agar plates exhibiting stronger lethality in acetic acid-fed conditions **(Figure 1B)**. These results are consistent with findings from Bruggeman et al. (2014) but differ slightly for oxyfluorfen, which may be due to experimental variations.

**Figure 1:**
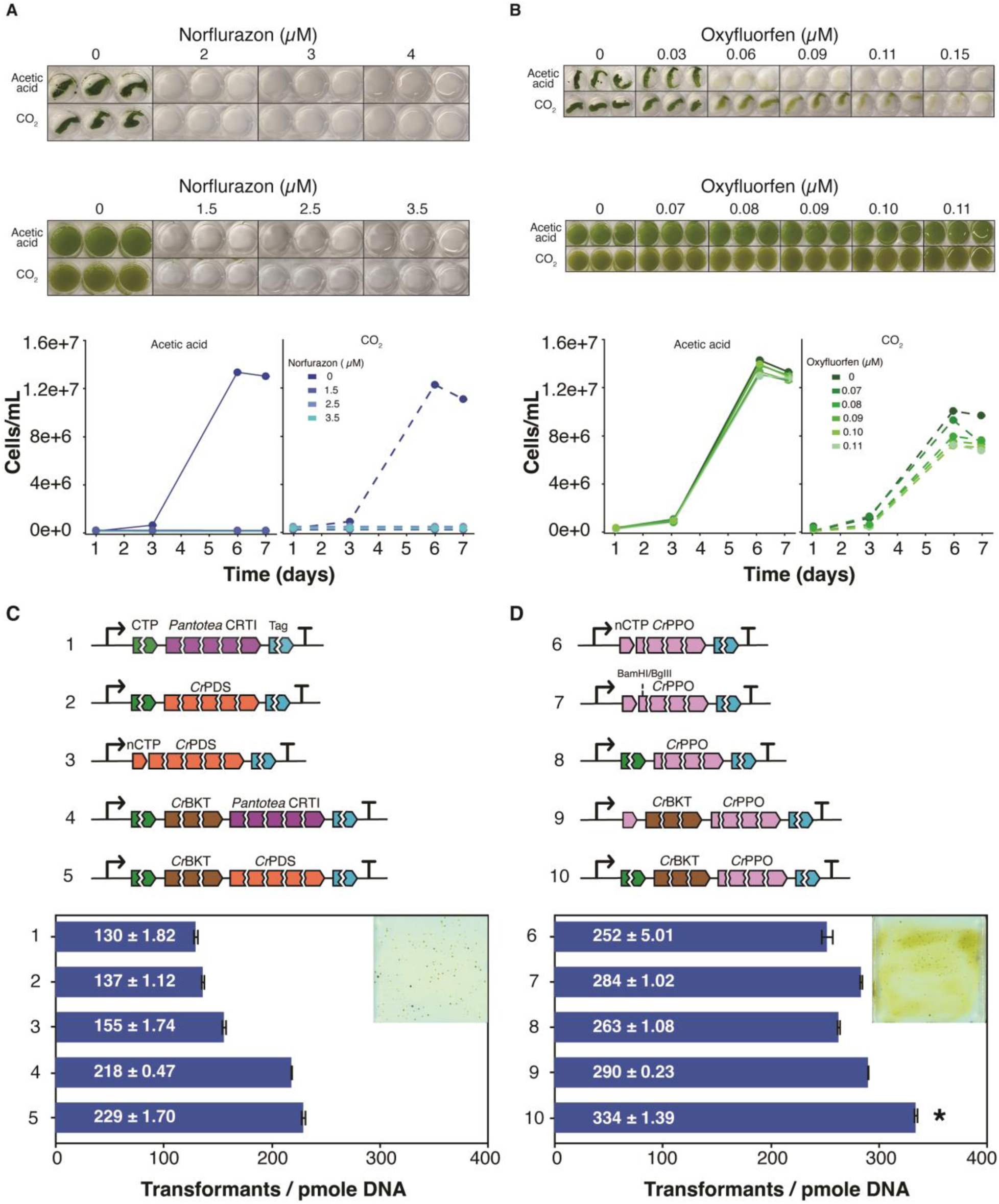
**Sensitivity of C. reinhardtii to norflurazon and oxyfluorfen herbicides and efficacy of resistance markers for transformant selection**. **A,B** – determination of norflurazon or oxyfluorfen concentrations that inhibit *C. reinhardtii* growth in liquid and agar solidified medium with either acetic acid or CO_2_ as a carbon source. Representative photographs and cell density of liquid cultures measured daily. **C,D** – plasmids used to confer resistance to each herbicide, all genes are synthetic variants with codon optimization and intron spreading. CTP – PsaD chloroplast targeting peptide, Pantotea CRTI – phytoene desaturase CrtI from *P. ananatis*, CrPDS – *C. reinhardtii* phytoene desaturase with R268T amino acid substitution, nCTP – native chloroplast targeting peptide, CrBKT – *C. reinhardtii* beta carotene ketolase, CrPPO – *C. reinhardtii* protox with V389M substitution. Complete annotated plasmid sequences are available in Supplemental Data 1. Below panels show transformation efficiency for each plasmid as the average of three biological replicates (n=3). Error bars represent standard deviation from mean. Photo inlays show transformant colonies generated and background cell debris. Asterisk (*) indicates the only construct which generated brown cells indicative of ketocarotenoid formation.

### 3.2 Synthetic phytoene desaturase and protox genes as selection markers

To test the efficiency of both herbicides as selection markers, ten different plasmids containing synthetic codon-optimized and intron-containing transgene sequence variations with known mutations that confer resistance were generated for phytoene desaturase (PDS) or protoporphyrinogen oxidase (PPO, ‘protox’) (**Figure 1C,D; Plasmids 1-10**). Norflurazon resistance was tested with the *P. ananatis* CrtI or the *Cr*PDS(R268T). Oxyfluorfen resistance was tested with the *Cr*PPO(V389M). Variants of each, using either the native chloroplast target peptide (CTP) of the gene, or the photosystem I reaction center subunit II (PsaD) CTP were all driven by the PsaD promoter. Fusions were also attempted with the *Cr*BKT to each gene to emulate previous fusions of this gene to the *E. coli aadA* that achieved selection and carotenoid ketolation phenotype in a single transformation (Figure 1C,D) (Cazzaniga et al., 2022).

Of the three variant PDS plasmids, Plasmid 3, containing the native CTP of this gene exhibited the highest transformation efficiency of 155±1.74 colonies / µg DNA. Although *Cr*BKT fusions to both PDS variants resulted in higher transformation efficiencies, neither resulted in red coloration. Norflurazon exhibited effective selection, with little cellular debris background on the agar plate **(Figure 1C, photo inlay)**. Oxyfluorfen resistance resulted in more colonies for all *Cr*PPO(V389M) variant plasmids than on norflurazon (**Figure 1D, Plasmids 6-10)**. In Plasmid 7, nucleotides were added after the predicted CTP to facilitate replacement of the *Cr*PPO(V389M) with the PsaD CTP (Plasmid 8). Use of the PsaD CTP and *Cr*BKT fusion to *Cr*PPO(V389M) in plasmid 10 led to 334±1.4 colonies / µg DNA and was also the only construct that led to brown coloration caused by ketocarotenoid formation **(Figure 1D, asterisk)**. Oxyfluorfen led to higher background cell debris and incomplete death; however, resistant colonies could be picked from the background **(Figure 1D, photo inlay)**. Plasmids 3 and 10 were then chosen for further use as herbicide-resistant selection markers to engineer *C. reinhardtii* for isoprene production.

### 3.3 Herbicide-resistance selection markers facilitate engineering isoprene production

We previously reported the generation of a strain of *C. reinhardtii*, which expressed three heterologous transgenes to facilitate metabolic channeling of carbon towards volatile isoprene production. The *Ib*IspS isoprene synthase, the *Cr*BKT beta carotene ketolase, and *Sc*IDI isomerase were expressed from the nuclear genome of the alga through three consecutive transformation and selection steps on paromomycin, spectinomycin, and zeocin selection, respectively (Yahya et al., 2023). Expression cassettes were developed here to enable *Ib*IspS and *Sc*IDI overexpression using the *Cr*PDS(R268T) (Plasmid 3) and *Cr*BKT-*Cr*PPO(V389M) fusion (Plasmid 10) herbicide resistance markers (Yahya et al., 2023). Six new plasmids, in addition to Plasmid 10, were designed and cloned to express combinations of *Ib*IspS, *Sc*IDI, and *Cr*BKT (when not fused to *Cr*PPO(V389M)) fused to fluorescent reporter proteins using the two herbicide-resistant selection markers **(Plasmids 11-16, Figure 2A)**. In parallel, we determined that isoprene production was highest when these genes were expressed in independent cassettes rather than in fusion proteins with one another **(Plasmids 17-29, Supplemental Figure 3).** We also determined that further modifications of the carotenoid pathway through co-expression of *C. reinhardtii* beta carotene hydroxylase (*Cr*CHYB) did not change isoprene production beyond that observed from the perturbation of expressing *Cr*BKT **(Supplemental Figure 4)**.

**Figure 2:**
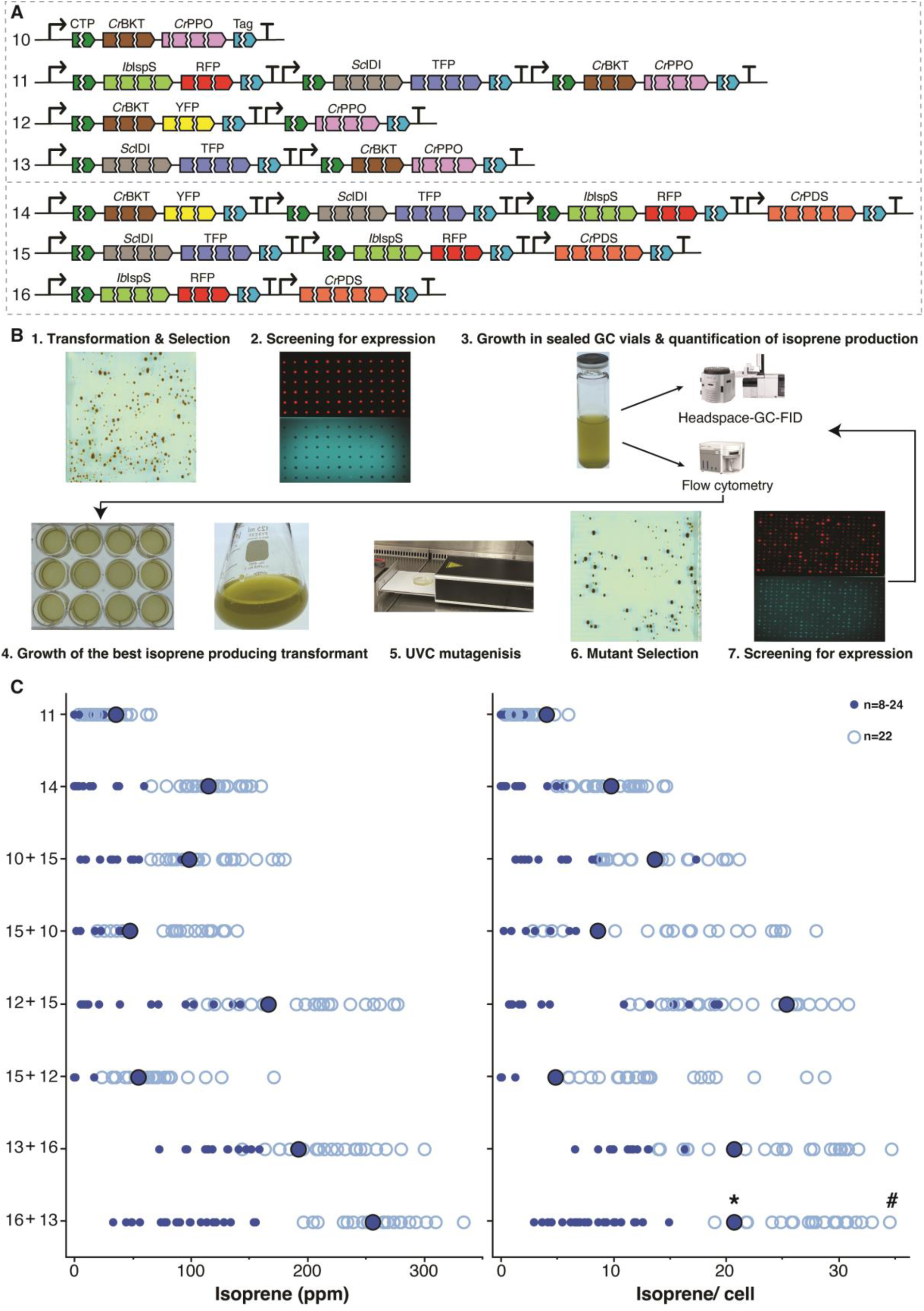
**Herbicide resistance genes and UV-C mutagenesis to select expression of isoprene metabolic engineering genes of interest**. **A** – plasmids investigated for expression of enzymes that can drive isoprene bioproduction from *C. reinhardtii*. Labels as Figure 1, with IbIspS – I. batatas isoprene synthase, RFP – mScarlet fluorescent protein, ScIDI – *S. cerevisiae* isomerase, TFP – mTFP fluorescent reporter, YFP – mVenus fluorescent reporter. Complete annotated plasmid sequences are available in Supplemental Data 1. **B** – workflow of transformant selection, isoprene quantification, subsequent UV-C mutagenesis, and further mutant isolation and isoprene production assessment. **C** – isoprene productivity measured in the headspace of 20 mL sealed vials for transformants (dark circles) and the mutants (light circles) derived from the most productive transformant strain for each group (black outlined dark circles). Each circle represents a single biological replicate for a population of picked transformants or mutants, as indicated. Asterisk and # symbols are individuals that were used in subsequent photobioreactor experiments.

These plasmids were transformed into UPN22 and fluorescence screening was used to identify fusion protein expression in isolated transformants as described in the Methods section. Correct molecular mass of each fusion protein was confirmed through SDS PAGE fluorescence **(Supplemental Figure 5)**. An attempt was made to achieve expression of all three target transgenes in a single plasmid (Plasmids 11 and 14), while plasmids 12, 13, 15, and 16 contained different combinations of *Ib*IspS, *Sc*IDI, *Cr*BKT and the two selection markers **(Figure 2A).** These were subsequently transformed in both sequence orders. After transformation and herbicide selection, all confirmed transformants, were grown in liquid medium in sealed GC vials with two replicates for each transformant to quantify isoprene production by day 6 of cultivation (**Figure 2B,C - dark blue circles**).

Plasmid 11 and plasmid 14 are larger (18 and 22kb respectively) compared to other plasmids (∼8- 17 kb) and contain all three GOI with either of the two selectable markers: *Cr*Protox(V389M) Plasmid 11 and *Cr*PDS(R268T) Plasmid 14. These two plasmids resulted in low isoprene yields **(Figure 2C)**. Although Plasmids 11 and 14 contain all three essential genes, few colonies could be obtained that expressed all the desired enzymes to reasonable levels. Position effects and plasmid shearing on transformation are known issues for large plasmids in nuclear transformation of *C. reinhardtii* (Lauersen, 2019; Perozeni & Baier, 2023; Zhang et al., 2021). The five other plasmids were designed for two rounds of transformation aimed at enabling selection of transformants with robust expression of each desired transgene to match those previously obtained with antibiotic selection (Yahya et al., 2023).

The best performing isoprene-producing strains were obtained from serial transformation and screening of plasmids 16+13 in either sequence order. Plasmid 16 has *Ib*IspS–RFP in one cassette, with *Cr*PDS(R268T) selection, and plasmid 13 has *Sc*IDI-mTFP in one cassette and *Cr*BKT- *Cr*Protox(V389M) for ketocarotenoid biosynthesis and selection, respectively. Those transformants generated with either order produced a maximum of ∼21 pg isoprene/cell **(Figure 2C, dark blue circles)**. These findings align with previous experiments using antibiotic resistance markers, which indicate that each enzyme performs optimally when positioned at the N-terminus of a fusion protein and when expressed separately in distinct cassettes (Yahya et al., 2023) **and Supplemental Figure 3**).

### 3.4 Isoprene production can be enhanced through further mutagenesis

Although isoprene production was achieved, overall isolation and expression of transformants with robust GOI expression was 34% lower with herbicide resistance markers than previously achieved through antibiotic selection (Yahya et al., 2023). Mutagenesis via UV-C exposure was then performed on the highest isoprene-producing transformants obtained from each plasmid combination **(Figure 2B)**. Colonies recovered on plates exposed to 30 sec of UV-C were selected for further screening. Independent colonies, hereafter ‘mutants’ were screened again for fluorescence signal from each fusion protein and subsequently grown in two biological replicates in GC vials for isoprene production assessment **(Figure 2B)**. All mutants are represented as light blue open circles in **Figure 2C**. Compared to the original transformation, a change in volumetric isoprene yield was observed in recovered mutants, with the majority improving from the original transformant. The best isoprene-producing mutants were obtained from strains transformed with Plasmids 16+13 in either order, each maximally producing ∼35 pg/cell, up to 334 mg/L **(Figure 2C)**. These results are comparable to those obtained with antibiotic selection (Yahya et al., 2023).

To further investigate whether observed increases in isoprene titers were caused by selection of transformants with increased transgene expression or background mutations leading to higher isoprenoid flux, in-gel SDS-PAGE fluorescence analysis was performed with the three best transformants of plasmid 16+13 and the three best mutants of the same plasmid **(Supplemental Figure 6**). Fluorescence intensity revealed higher heterologous protein expression levels for both *Ib*IspS-RFP and *Sc*IDI-mTFP in the mutants compared to the original transformants, correlating with the increased isoprene yields observed **(Supplemental Figure 6)**. This result confirms that the mutations likely result in selection of strains with increased transgene expression, leading to observed higher isoprene yields.

### 3.5 Isoprene is produced from engineered *C. reinhardtii* grown on CO_2_ as a carbon source

Up to this point, analysis of engineered isoprene production in *C. reinhardtii* was conducted through growth on acetic acid as a carbon source in sealed vials to facilitate high-throughput headspace analysis (see Methods). We decided to explore whether we could use yeast respiration as a source of CO_2_ to grow the alga in sealed vials. As *S. cerevisiae* cannot metabolize nitrate and *C. reinhardtii* cannot metabolize glucose, co-culture of these organisms in TAPhi-NO_3_ or TPhi-NO_3_ medium with glucose supplementation provided an opportunity to use yeast respiration of glucose to generate CO_2_, without it overtaking the algal culture **(Supplemental Figure 7)**. It was possible to conduct these co-cultivations with isoprene-producing *C. reinhardtii* strains and measure isoprene production in the headspace of sealed vials. Without acetic acid, algal cells were able to grow and showed a correlation between cell density and isoprene production, which followed initial glucose concentrations **(Supplemental Figure 7)**. We also tested the impact of ethanol produced by yeast on *C. reinhardtii,* which did not negatively affect algal growth or isoprene production in acetic acid-containing media and did not serve as a carbon source for the alga **(Supplemental Figure 8)**. The results may be of use to others wishing to test CO_2_-based carbon source growth for *C. reinhardtii* in sealed vessels.

### 3.6 Multi-port mass spectroscopy coupled to parallel photobioreactors enables real-time isoprene production analysis from CO_2_ as a carbon source

Metabolic engineering of algae aims at generating strains that could be used in larger-scale cultivation concepts, preferably to convert inorganic wastes into higher chemical order value. Algal photobioreactors use CO_2_ as the carbon source; hence, we were interested in testing the engineered alga’s ability to produce isoprene in photobioreactors instead of simply screening conditions. Recently, we reported on the possibility of coupling a multi-port inlet mass spectrometer to parallel photobioreactors (as illustrated in (Villegas-Valencia et al., 2025). Here, we used this setup to systematically test isoprene production from *C. reinhardtii* grown on CO_2_ as a sole carbon source in 400 mL cultivations.

### 3.7 Warmer temperatures and constant illumination improve isoprene production

Our previous results indicated that temperature had a larger impact on isoprene production than light intensity in sealed vials using acetic acid as a carbon source for growth (Yahya et al., 2023). With the ability to tightly regulate cultivation conditions in photobioreactors, we sought to test the impact of these abiotic factors on both culture growth performance and volatile isoprene production. The highest isoprene-producing transformant (prior to mutagenesis), carrying Plasmids 16+13, which yielded 256 mg isoprene/L in screening conditions **(Figure 2)**, was used in the following experiments. Two preliminary experiments assessed the impact of temperature and illumination cycles on isoprene production from *C. reinhardtii* grown on CO_2_. The first experiment tested temperatures (25 °C and 30 °C) and light regimes (constant light and a 16h:8h light:dark cycle). Continuous light at 30 °C resulted in the highest isoprene production, reaching 172 mg isoprene/L after 21 days. A temperature increase to 38°C at the end of cultivation led to a brief spike in isoprene production across all conditions **(Supplemental Figure 9A-C)**.

A second trial explored the effect of different temperatures (30 °C to 34 °C) under continuous light, with heating cycles where the temperature was briefly spiked to 38°C at different intervals **(Supplemental Figure 10)**. Consistent cultivation at 34 °C yielded the highest cumulative production, reaching ∼175 mg isoprene/L culture on day 8 **(Supplemental Figure 10F)**. Although overall production was lower with a base temperature of 30 °C, daily heating cycles to 38 °C improved isoprene yield over this baseline in all tested variations **(Supplemental Figure 10F)**. As the boiling point of Isoprene is at 34 °C, it is logical that temperature would significantly affect the measurable yields of isoprene, especially with gas flowthrough in photobioreactors.

### 3.8 Multi-parallel parameter testing reveals optimal temperature and illumination conditions

We next systematically tested a range of 35 different combinations of light intensities (300, 600, 900, 1200, and 2000 µE) and temperatures (30, 31, 32, 33, 34, 35, and 36 °C) using the multi-parallel photobioreactors to refine abiotic settings that would maximize cell growth and isoprene yield in batch CO_2_-fed cultivations **(Figure 3)**. The highest optical density (OD_740_) values were recorded between 31 and 33 °C. As light intensity increased to 1200 µE, OD740 values remained relatively high across these temperatures. Higher temperatures of 35 and 36 °C led to a notable reduction in OD_740_ values, especially at 300 µE, where the lowest OD_740_ (1.9) was observed at 36 °C **(Figure 3A).** Overall, temperatures of 33 and 34 °C consistently exhibited the highest isoprene levels across all light intensities, likely as these are near the growth optimum of *C. reinhardtii* (**Figure 3B**). The optimal condition was identified as 33 °C and 900 µE, which provided the highest daily isoprene yield normalized to optical density.

**Figure 3:**
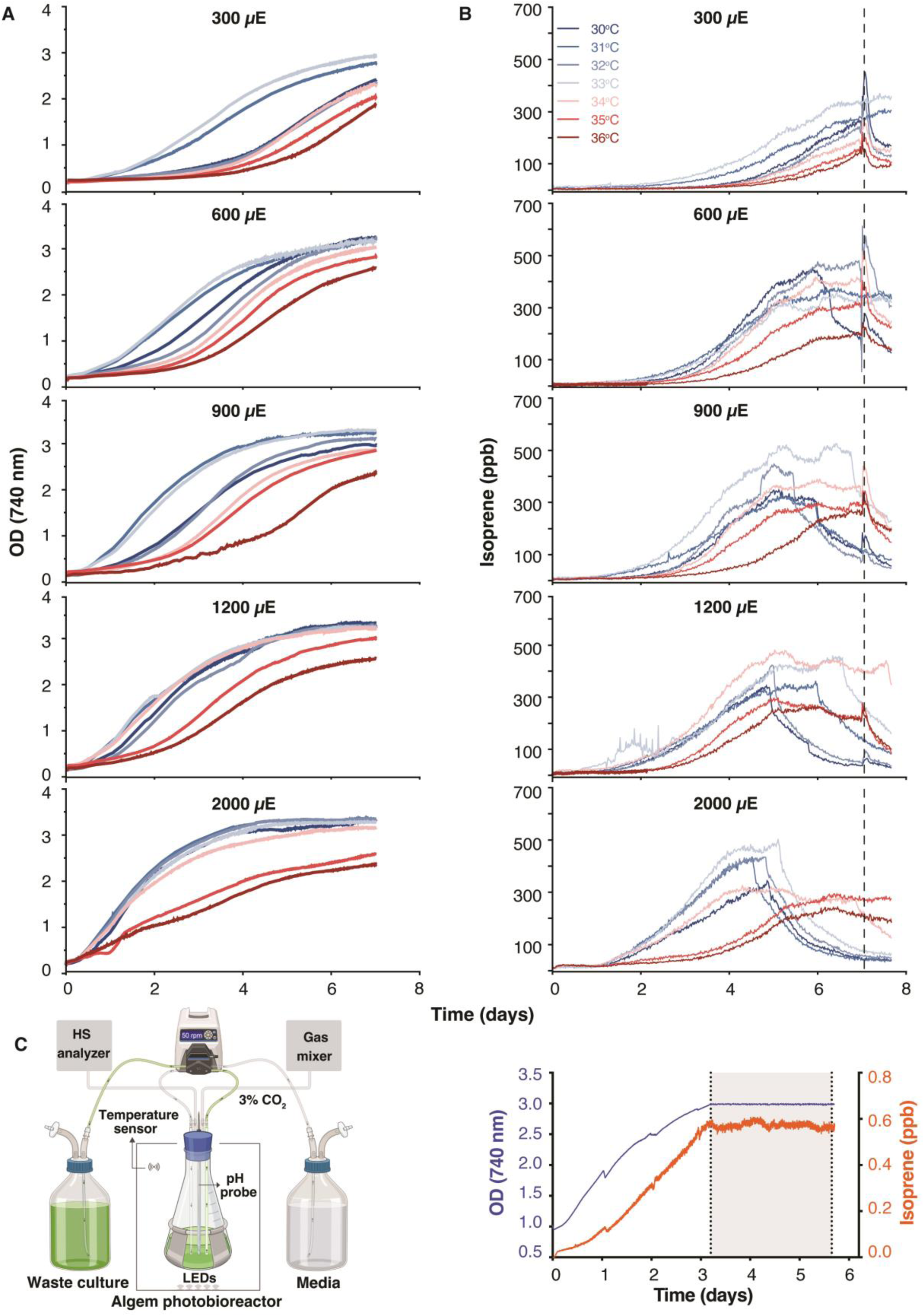
Growth performance and in-line analysis of volatile isoprene in off-gas from 400 mL photobioreactors growing engineered *C. reinhardtii*. **A,B** – optical density (OD) at 740 nm measured in 15 min intervals and direct off-gas isoprene readings of best-producing mutant cultivated in separate parallel 400 mL photobioreactors subjected to 35 different combinations of continuous light and temperature conditions coupled to the multi-port inlet real-time headspace gas analysis system. **C** – schematic of a turbidostat photobioreactor cultivation setup and the measured optical density and live isoprene readings during establishment and continuous production when maintained at an OD(740nm) of 3.0 by the turbidostat module for 3 days (grayed portion). Section C was created with BiorRender.com.

### 3.9 Continuous isoprene production in turbidostat mode

Using these optimal settings, a culture of mutant from Plasmids 16+13 was cultivated in turbidostat mode to determine if isoprene production could be sustained at a constant rate **(Figure 3C)**. Batch preculture was grown using 33 °C and 900 µE as above until OD 3.0 was reached. Upon activation of the turbidostat pumps, the culture density could be maintained in a tight window, and live isoprene production also exhibited steady readings just under 0.6 ppb. Consistent cultivation in this mode lasted for 36 h and averaged 51.2 ± 0.7 mg isoprene / L / day (**Figure 3C**). Given the density of isoprene (681 g/L), at this production rate, it would require ∼13,300 L of algal culture to yield 1 L of isoprene/day. Although likely not an economically relevant yield on which to build an isoprene production bioprocess, this represents a separately harvestable co-product from the algal biomass. It should also be noted that these strains produce a mixture of ketocarotenoids, which can increase the value of the algal biomass. One goal of engineering co-products in algal hosts could be their use in wastewater treatment. We next explored whether we could concomitantly generate isoprene while removing contaminating nitrogen and phosphorous in effluents from an anaerobic membrane bioreactor (AnMBR) wastewater treatment process.

### 3.10 Nutrient limitation affects isoprene production in repeated batch cultivation using wastewater as culture medium

The best mutant strain from the previous experiment, which produced 334 mg isoprene / L in screening conditions, was selected to evaluate whether *C. reinhardtii* grown on wastewater could yield isoprene, reduce contamination of inorganics, and produce biomass. To this end, repeated batch cultivations were conducted using two different growth media: Replete medium (T2Phi- NO₃) and AnMBR effluent. The treatments included T2Phi-NO₃, AnMBR WW raw, AnMBR WW + Trace Nutrients from T2Phi-NO₃, AnMBR WW + NH₄⁺ at the concentration of nitrogen (as ammonium) in T2Phi-NO₃, and AnMBR WW + PO₃³⁻ at the concentration of T2Phi-NO₃. Cultures were grown for 8 days under continuous illumination (900 μE) at 33°C, with constant CO₂ sparging (3% CO₂ at 25 mL/min). Every 48 h, 350 mL from the 400 mL culture was replaced with fresh medium according to the respective treatment. Growth was monitored via OD₇₄₀ measurements, and biomass concentrations were calculated using a standard curve. Isoprene production was quantified in real-time using the headspace gas analysis system above.

Biomass concentrations were similar across all treatments, with maximum values of 1 g/L for T2Phi-NO₃, 0.9 g/L for AnMBR WW, 0.8 g/L for WW + Trace Nutrients, 0.8 g/L for WW + NH₄⁺, and 0.8 g/L for WW + PO₃³⁻. No significant differences in maximum biomass concentrations were observed between treatments over the 8-d cultivation period **(Figure 4A)**. Isoprene production was highest and most stable in the T2Phi-NO₃ treatment, maintaining consistent levels throughout the experiment and reaching a peak of 642 ppb. In contrast, although biomass was consistently formed, the AnMBR WW treatment exhibited a gradual decline in isoprene production after each successive dilution, starting at 579 ppb and decreasing to 446 ppb after 96 h, 373 ppb at 144 h, and 302 ppb at 192 h. This pattern suggests that the untreated AnMBR WW lacks a crucial trace element required for isoprene metabolism. The addition of trace nutrients (WW + Trace Nutrients) led to more stable isoprene production (395–359 ppb) across repetitions **(Figure 4)**, suggesting partial alleviation of nutrient limitations. Overall production in this condition remained lower than in T2Phi-NO₃, likely due to insufficient macronutrients such as nitrogen and phosphorus. Supplementation with ammonium (NH₄⁺) in the WW + NH₄⁺ treatment further increased isoprene production to 404 ppb, likely due to improved nitrogen availability, while phosphite (PO₃³⁻) addition in the WW + Phosphite treatment led to a peak of 340 ppb **(Figure 4A)**. The results are important for setting parameters if a WW treatment process was established with engineered algae.

**Figure 4:**
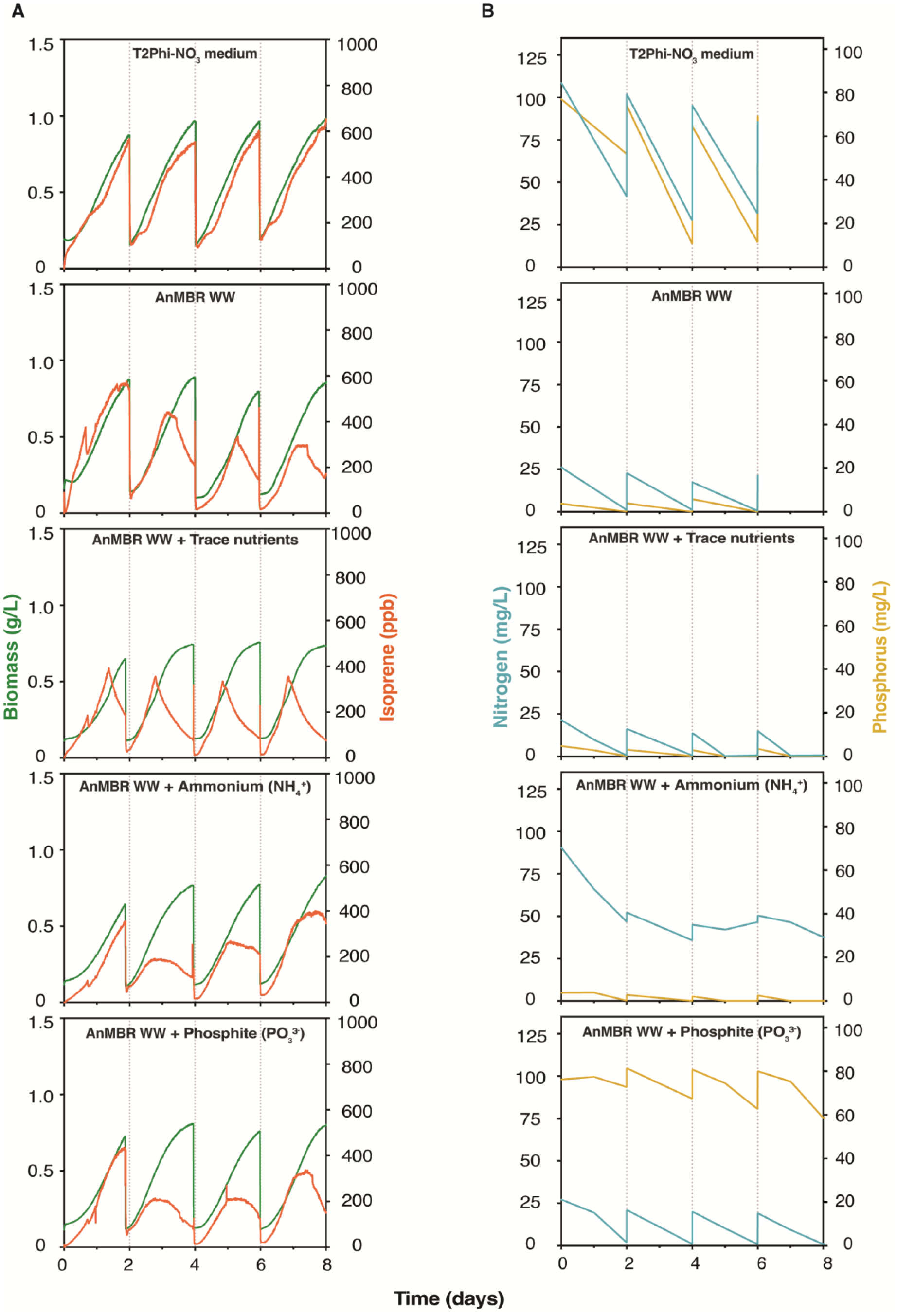
Analysis of culture growth and isoprene production in the off-gas of parallel photobioreactor cultivations of engineered *C. reinhardtii* grown in replete medium or anaerobic membrane bioreactor (AnMBR) effluent. **A** – Estimated biomass (g/L) based on OD_(740nm)_ measurements (green) and direct off-gas isoprene (orange) readings of batch cultivations of best-producing mutant. Every 48 hours, 350 mL of the 400 mL culture volume was harvested and replaced with fresh medium. **B** – Quantified total nitrogen, ammonium or nitrate, (in blue), and phosphorous (in yellow) in culture medium, measured every 24 hours during each batch cycle showing depletion of N and P from AnMBR wastewater (WW).

## 4 Conclusions

*C. reinhardtii* has emerged as a promising photosynthetic host for heterologous metabolite production. In this work, we successfully used the herbicides norflurazon and oxyfluorfen, alongside the endogenous but optimized resistance genes *Cr*PDS(R268T) and *Cr*PPO(V389M), as selection markers for nuclear transformation in *C. reinhardtii*. These genes facilitated the overexpression of other genes of interest for metabolic engineering in separate cassettes. Although transgene expression under herbicide selection was less robust compared to previous reports with antibiotic selection, this limitation could be addressed through further UV-C mutagenesis, which allowed selection of mutants that more robustly expressed target GOIs. Here, coupling in-line headspace analysis to photobioreactors allowed continuous monitoring of volatile isoprene production from algal cultures grown on CO₂ as the sole carbon source. The capacity for multi-parallel cultivations and real-time monitoring enabled precise parameter definition to improve process outputs. The potential of *C. reinhardtii* to grow and produce isoprene using AnMBR effluent as a nutrient source supports future waste-stream valorization concepts, where engineered algae could both clean effluents and generate valuable co-products. The accumulation of ketocarotenoids in the biomass alongside isoprene production enhances the biomass value, making *C. reinhardtii* an attractive candidate for bio-refinery processes that combine multiple product outputs.

## 6 Author Contributions (CReDit)

**Razan Z. Yahya:** Conceptualization, Microbial cultivations, Experimental data collection, Data curation, Data analysis, Visualization, Writing – original draft

**Sebastian Overmans:** Conceptualization, Experimental data collection, Data curation, Data analysis, Visualization, Writing – review and editing

**Gordon B. Wellman:** Conceptualization, Plasmid design, Data curation, Data analysis, Writing –review and editing

**Kyle J. Lauersen:** Conceptualization, Funding acquisition, Project administration, Resources, Writing – original draft

## 7 Conflict of Interest

The authors declare that they have no conflict of interest.

## Supporting information

Plasmid Sequences

Data for all Figures

## Acknowledgements

The authors are grateful to Prof. Pascal Saikaly and Dr. Krishna Katuri for providing effluent from their AnMBR treatment process. KJL acknowledges baseline research funding from KAUST and ASEPC 106 funding of the Hiden HPR 20 system. The authors acknowledge additional financial support from a research grant with Saudi International Petrochemical Company (SIPCHEM).

## 9 Supplemental Figures

**Supplemental Figure 1: Isoprene standard curves in photobioreactor off-gas.** Generation of isoprene standard curve from the off-gas of 400 mL photobioreactors using the multi-port inlet real-time headspace gas analysis system, including a triple filter mass spectrometer (Hiden Analytical HPR-20 R&D, UK). **A** – observed isoprene peaks, from which the area under the curve was calculated. **B** – representation of observed mass spectrometer readings against the quantity of loaded isoprene into each bioreactor unit. Gas flow rates as described in the Material and Methods section.

**Supplemental Figure 2: Algem Turbidostat pumping volume standard curve.** Measured volumes of flow rates plotted against the pump speed setting on Algenuity, Algem photobioreactor turbidostat systems. The curve was used to set flow rates for turbidostat experiments.

**Supplemental Figure 3: Isoprene production in *C. reinhardtii* transformants of various plasmid constructs.** Isoprene in the headspace was measured from acetic acid-fed grown algal cultures in 20 mL sealed vials for screening. (A) Direct headspace isoprene measurement, (B) normalized Isoprene production per cell, and (c) normalized isoprene production per cell per YFP fluorescence of multiple independent transformants as indicated (n= 3–6). Spec – *E. coli aadA* spectinomycin resistance gene, YFP – mVenus fluorescent protein, *Ib*IspS – *I. batatas* isoprene synthase, *Sc*IDI – *S. cerevisiae* isomerase, *Cr*BKT – *C. reinhardtii* beta carotene ketolase. All constructs are driven by the PsaD promoter and use its plastid targeting peptide. Complete annotated sequences are presented in Supplemental File 1.

**Supplemental Figure 4: Comparison of isoprene yields in transformants with beta carotene hydroxylase and/or beta carotene ketolase. A** – plasmids used, *C. reinhardtii* beta carotene hydroxylase (*Cr*CHYb) and mTFP fluorescent protein are shown here for the first time in plasmids maps. Complete annotated sequences are presented in Supplemental File 1. **B** – SDS PAGE in gel fluorescence of signals from RFP and YFP fusions to target transgenes, with Western blotting of the CrBKT-aadA fusion (anti-aadA from Agrisera). **C,D** – isoprene yield in the headspace of transformants of indicated plasmid combinations grown in 20 mL seal vials on acetic acid.

**Supplemental Figure 5: SDS PAGE and in-gel fluorescence for molecular mass determination of expressed target fusion proteins**. The signal of each target recombinant protein-FP fusion from the indicated plasmids was determined to confirm whole-length expression.

**Supplemental Figure 6: SDS PAGE and in-gel fluorescence for expression comparison after UV-C mutagenesis**. The signal of each recombinant protein-FP fusion from transformants was determined before and after UV-C mutagenesis and further selection. Cell concentrations of transformants and selected mutants were normalized prior to protein extraction.

**Supplemental Figure 7: Co-culture of *C. reinhardtii* with *S. cerevisiae* enables CO_2_-based growth in sealed vials for isoprene production analysis. A** – photograph of co-culture vials and experimental conditions, Glu – glucose. **B,C** – Cell density of algae and yeast grown with acetic acid or without, respectively. Measured isoprene and CO_2_ concentration observed on days 6, 7, and 8 of growth in headspace of vials.

**Supplemental Figure 8: Effect of ethanol addition to *C. reinhardtii* culture grown with or without acetic acid in sealed vials.** Left – cultures with acetic acid, right – cultures without acetic acid. **A,B** – isoprene measured in the headspace. **C,D** – cell densities measured. **E,F** isoprene measured. **G,H** – ethanol measured in the headspace.

**Supplemental Figure 9: Pre-test of isoprene yields in 400 mL photobioreactors at different culture temperatures and illumination regimes. A** – plotted live readings of off-gas isoprene concentrations, **B** – cumulative isoprene concentrations, and **C** - daily cumulative isoprene concentrations measured using the multi-port inlet real-time headspace gas analysis system.

**Supplemental Figure 10: Pre-test of isoprene yields in 400 mL photobioreactors at different culture temperatures and heating regimes. A** – off-gas isoprene measured from cultures cultivated at three different temperatures with constant illumination. **B** – isoprene measured from cultures cultivated at 30 °C with three different regimes of periodic heating spikes to 38 °C. **C** – optical density measured from cultures cultivated at three different temperatures with constant illumination. **D** – optical density measured from cultures cultivated at 30 °C with three different regimes of periodic heating spikes to 38 °C. **E** – periodic heating cycles for cultures featured in B, D. **F** – cumulative isoprene measured using the multi-port inlet real-time headspace gas analysis system.

## 10 Supplemental Data Files

**Supplemental Data 1: Plasmid Sequences.** This file contains the annotated sequences in .gb format for all plasmids used in this work. It is a .txt file which should be opened with any plasmid editing software to generate annotated maps of 33 separate plasmids.

**Supplemental Data 2: Data for all figures presented in this work.** This file contains the raw data used in creation of all presented findings in this work. Each Figure’s data is separated by a unique sheet in the data file.Figures

## Notes

### Competing Interest Statement

The authors have declared no competing interest.

