## Supplemental Figures for "Engineered isoprene production from *Chlamydomonas reinhardtii* using herbicide selection markers and CO_2_-fed cultivation optimization through multi-parallel photobioreactor headspace gas analysis"

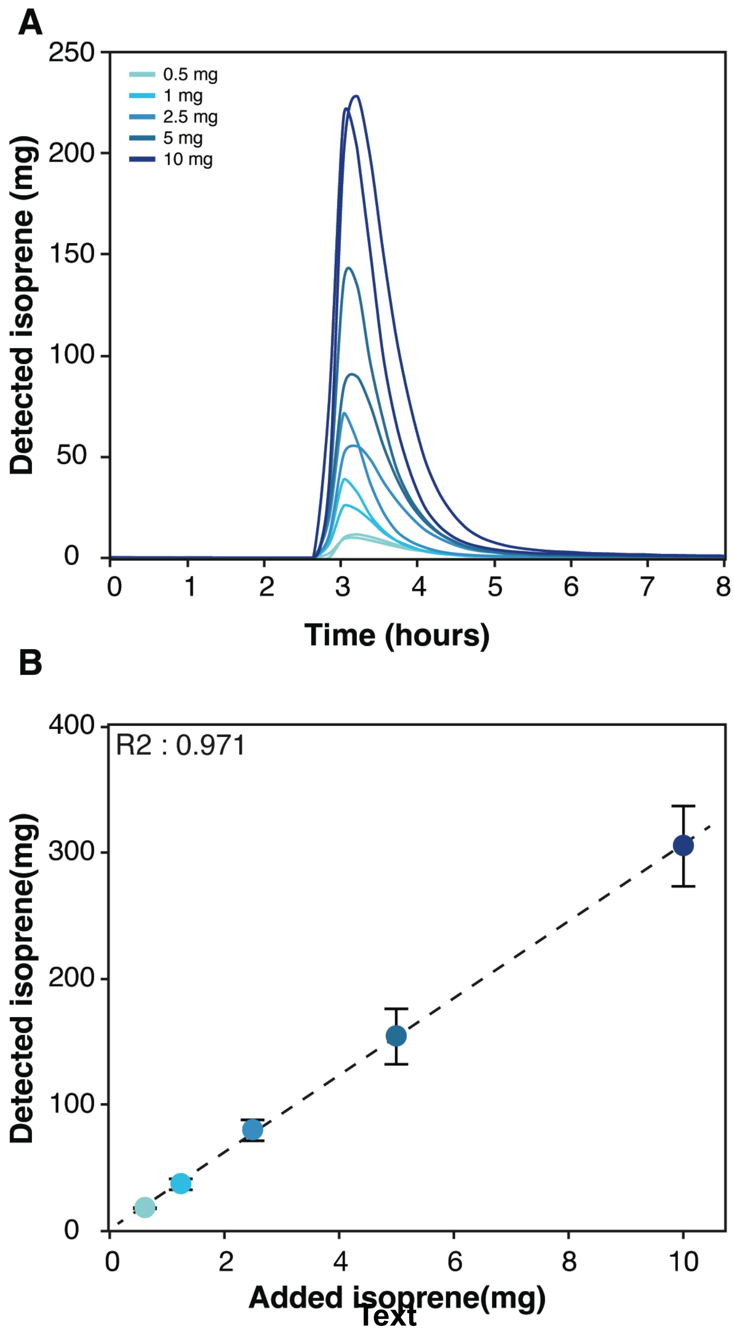


**Supplemental Figure 1**

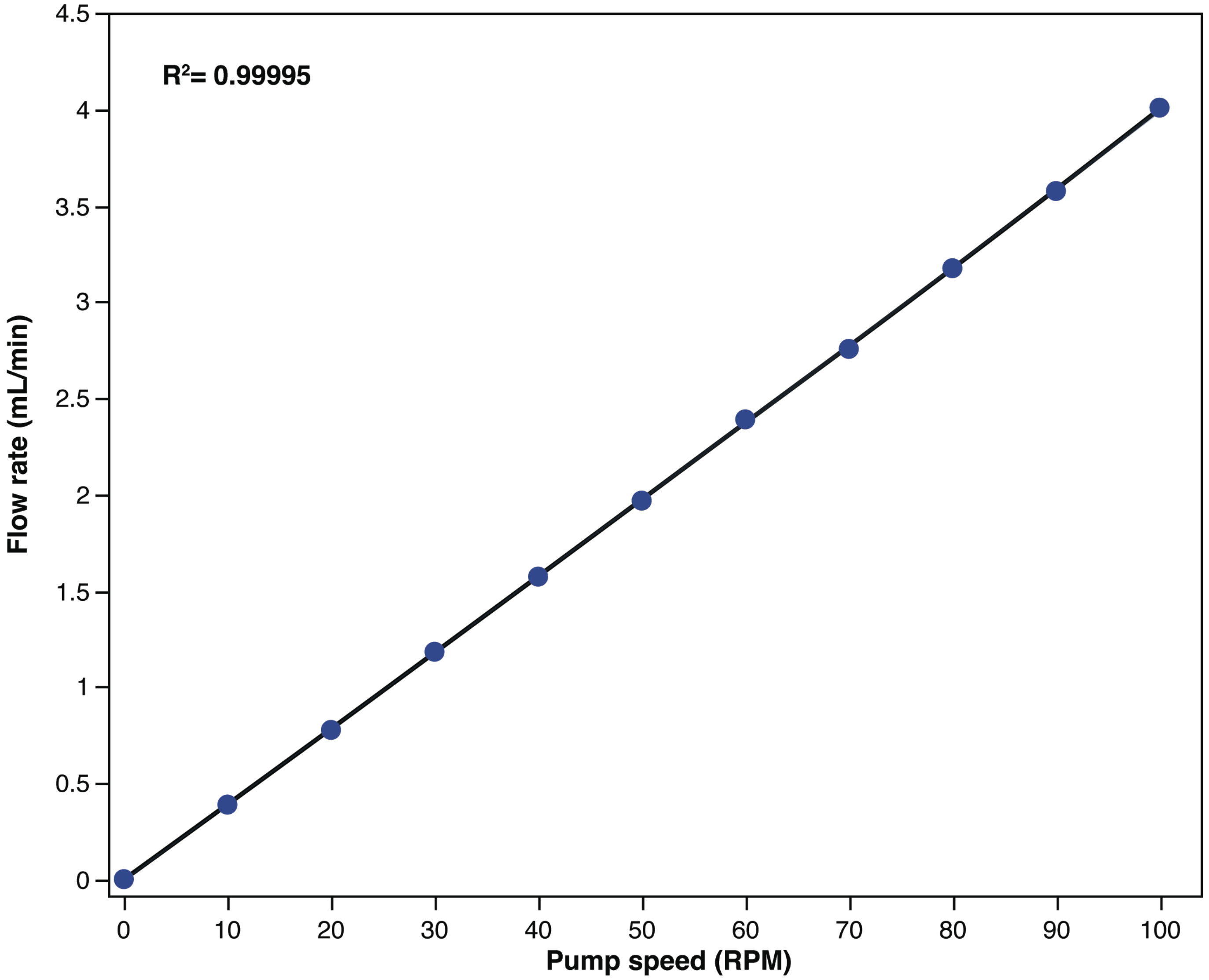


**Supplemental Figure 2**

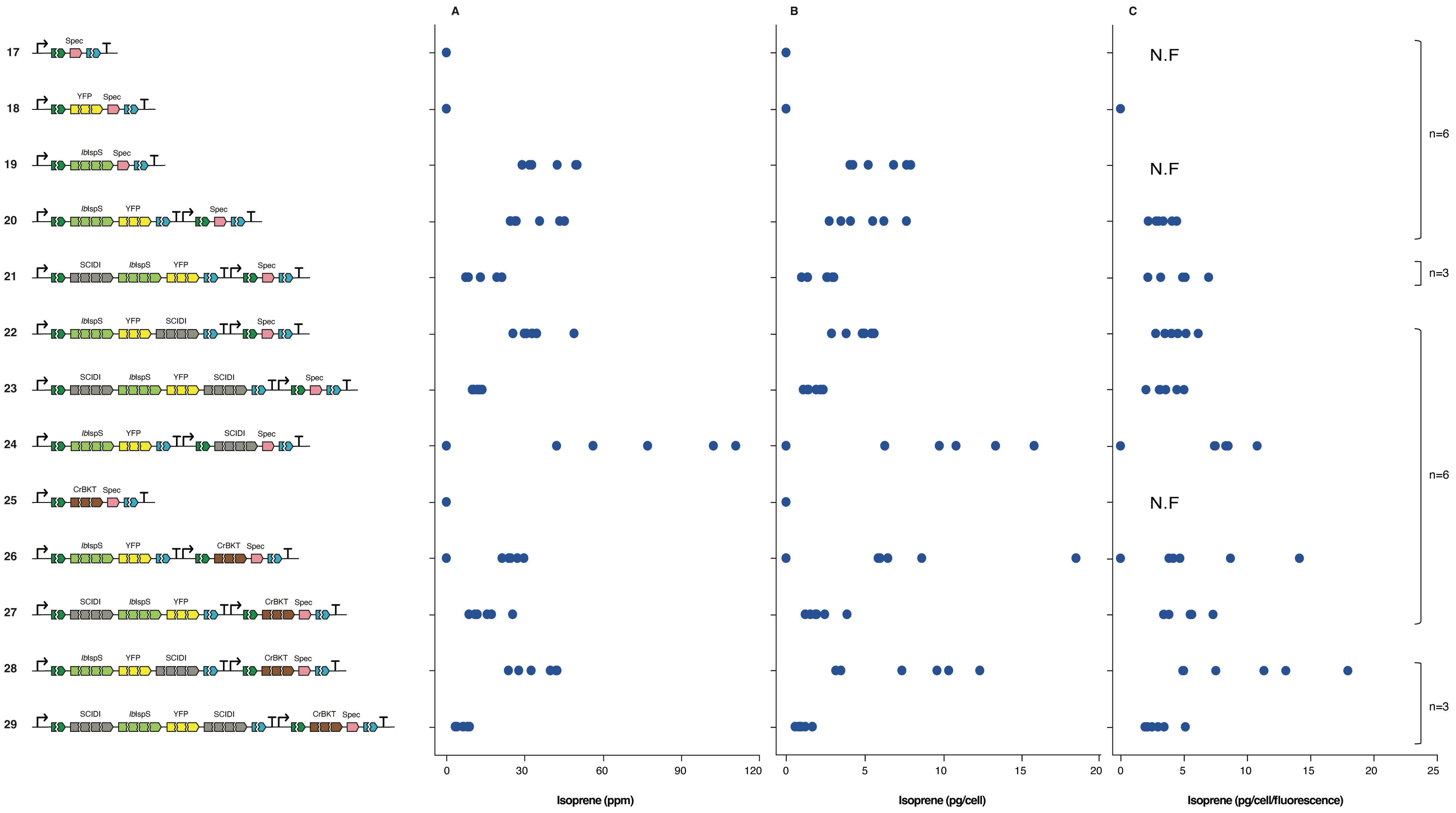


**Supplemental Figure 3**

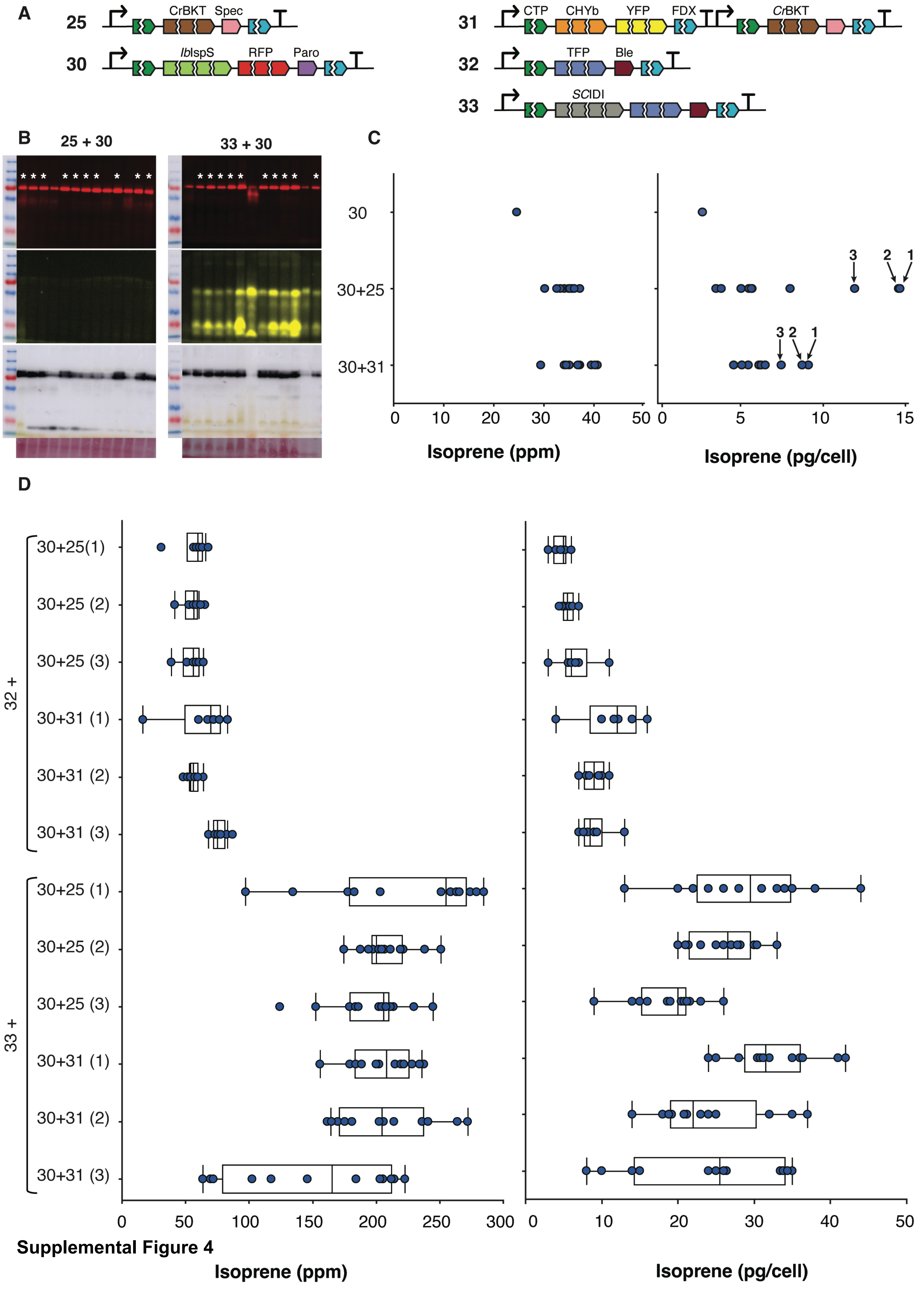


Supplemental Figure 4

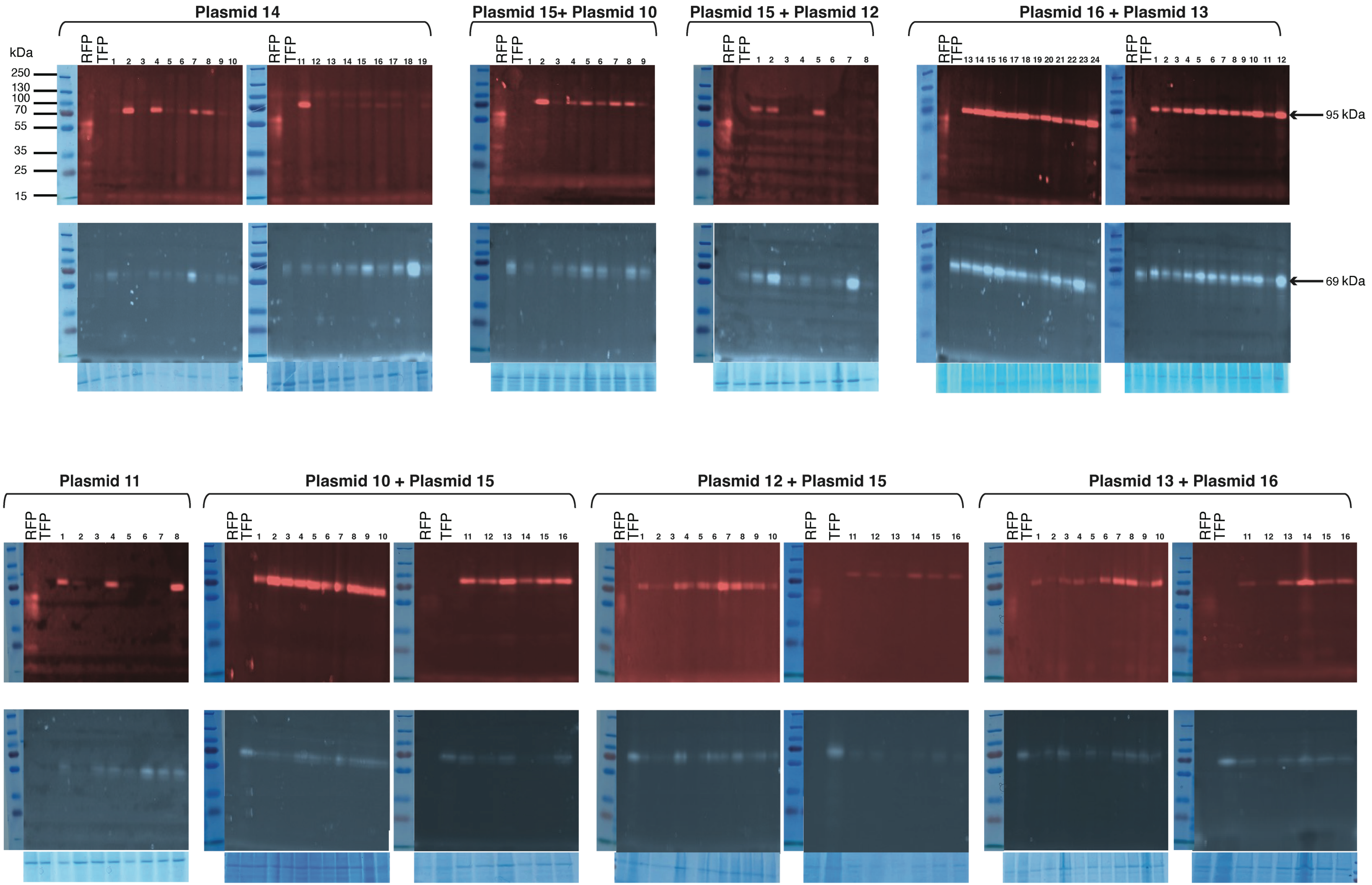


**Supplemental Figure 5**

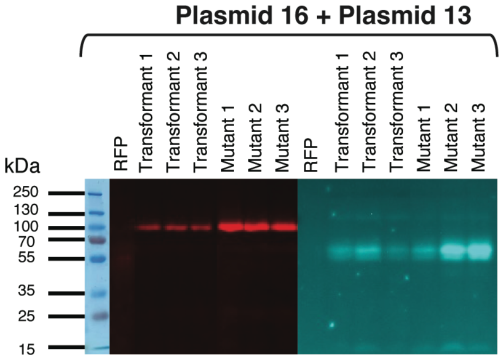


**Supplemental Figure 6**

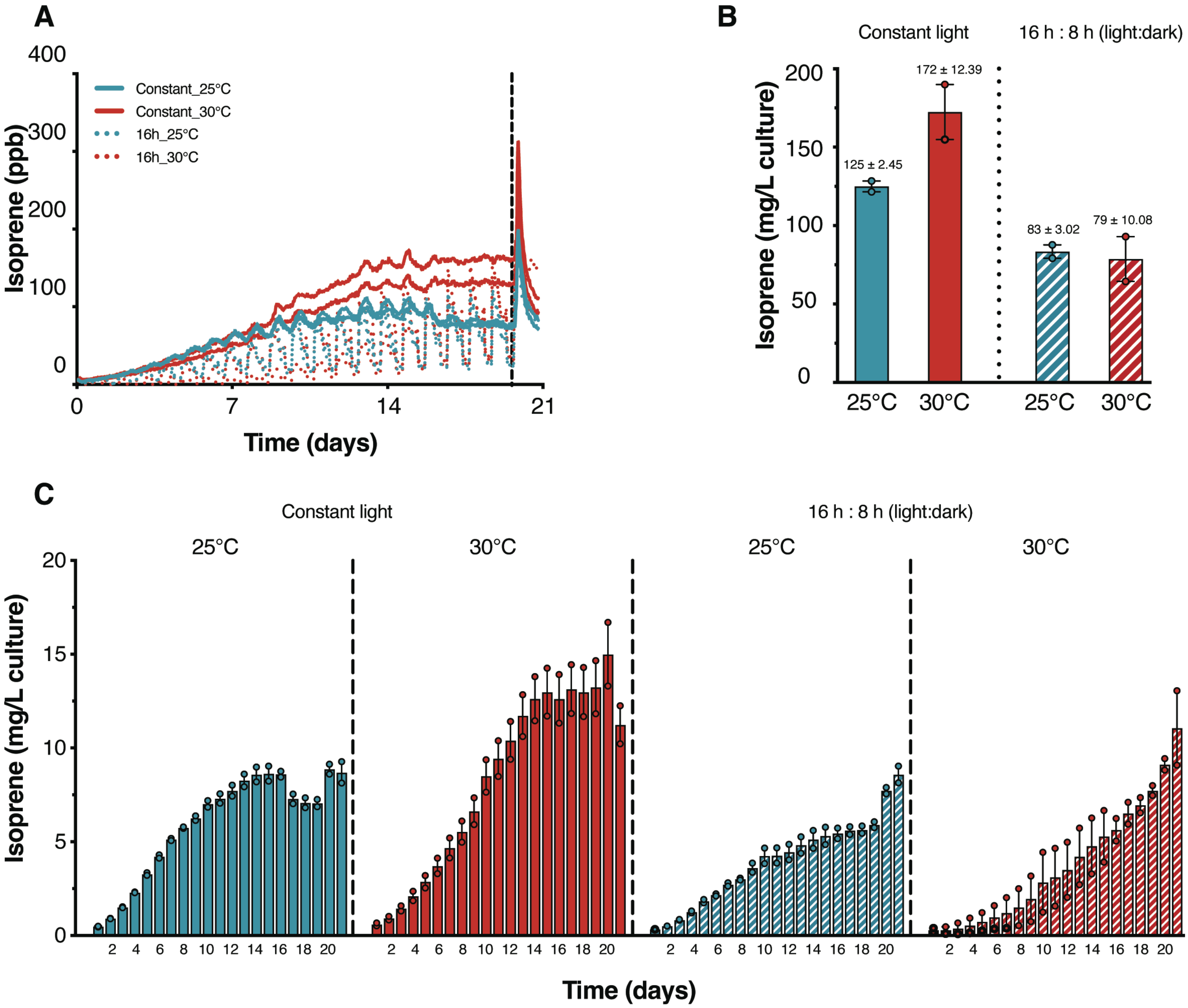


**Supplemental Figure 7**

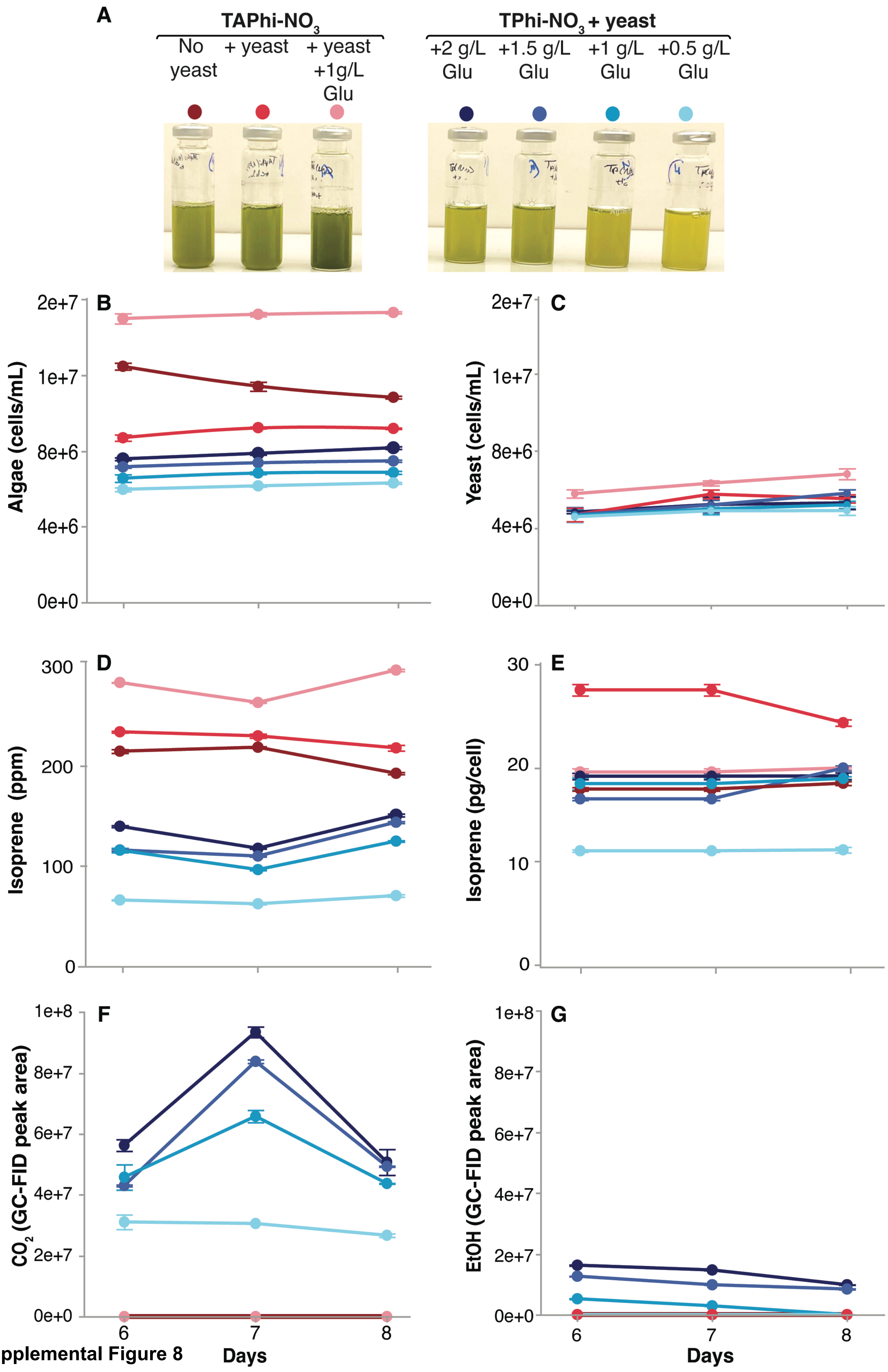


Supplemental Figure 8

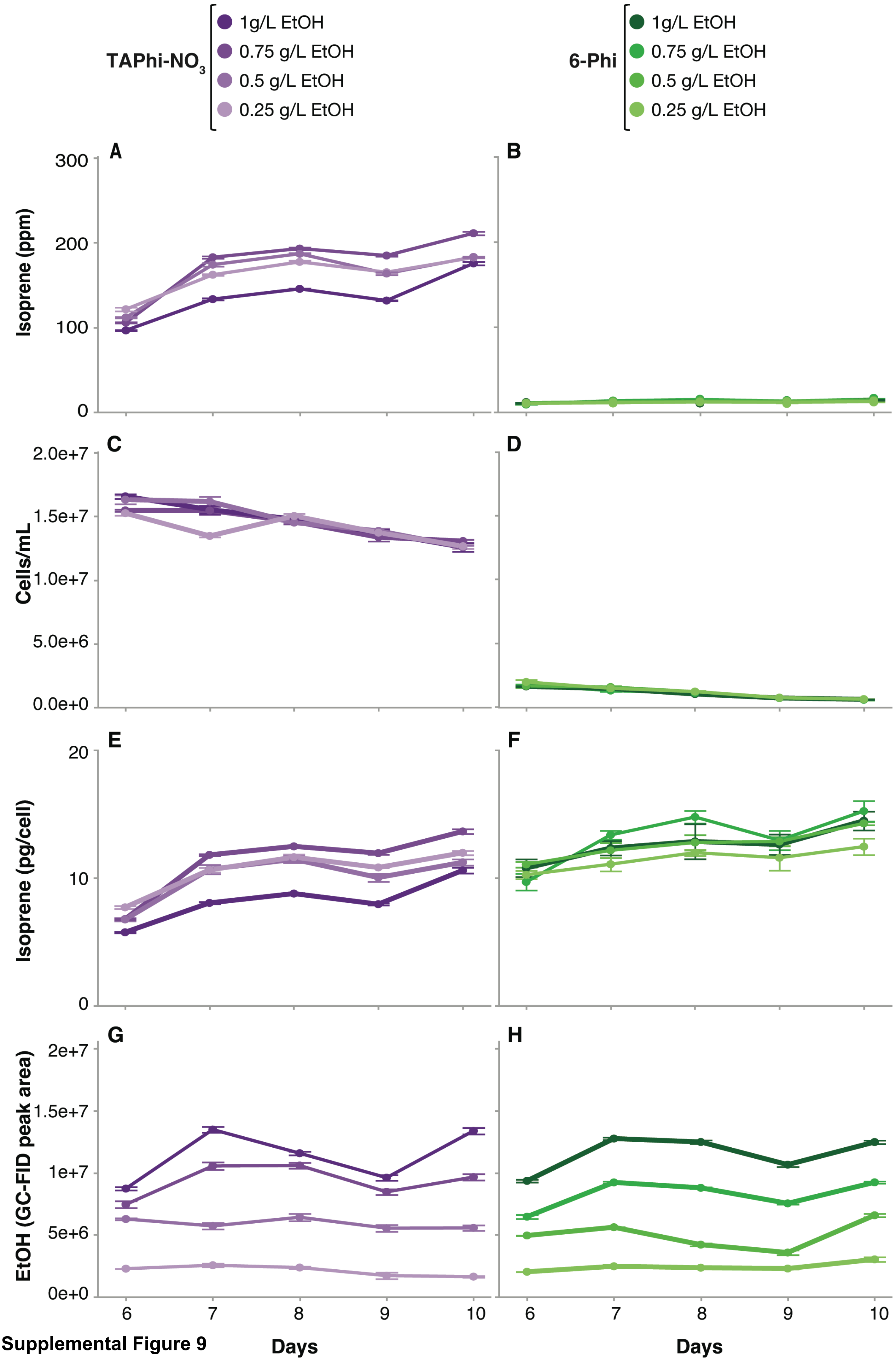


Supplemental Figure 9

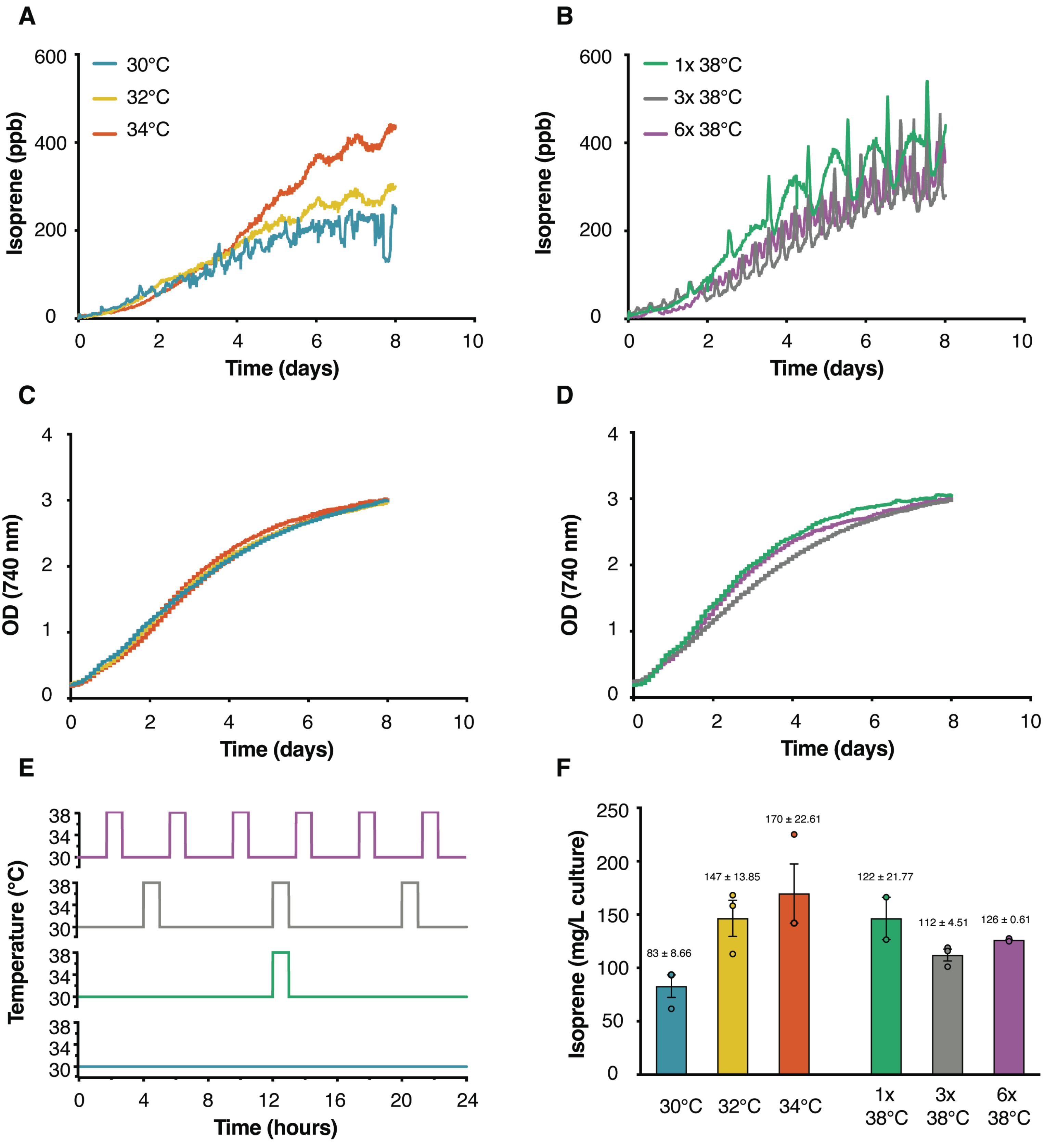


**Supplemental Figure 10**
